# Genome-wide association uncovers the genetic architecture of tradeoff between flowering date and yield components in sesame

**DOI:** 10.1101/2021.04.22.440889

**Authors:** Idan Sabag, Gota Morota, Zvi Peleg

## Abstract

Unrevealing the genetic makeup of crop morpho-agronomic traits is essential for improving yield quality and sustainability. Sesame (*Sesamum indicum* L.), one of the oldest oil-crops in the world, which despite its economical and agricultural importance, is an ‘orphan crop-plant’ that undergone limited modern selection, thus, preserving wide genetic diversity. Here we harnessed this natural variation in a newly developed sesame panel (SCHUJI) to perform genome-wide association studies for morpho-agronomic traits under the Mediterranean climate conditions. Field-based phenotyping of the SCHUJI panel across two seasons exposed wide phenotypic variation for all traits. Using 20,294 single-nucleotide polymorphism markers, we detected 50 genomic signals associated with these traits. Major genomic region on LG2 was associated with flowering date and yield-related traits, exemplified the key role of the flowering date on productivity. Our results shed light on the genetic architecture of flowering date and its interaction with yield components in sesame and may serve as a basis for future sesame breeding programs in the Mediterranean basin.

## 1. INTRODUCTION

Sesame (*Sesamum indicum* L.), one of the oldest oil-crops in the world, was domesticated about 5,500 years ago from *Sesamum indicum* subsp. *malabaricum* (Bedigian, 2015) in the Indian sub-continent. Sesame is an annual diploid (2n=2x=26) species, which belongs to the *Sesamum* genus from the *Pedaliaceae* family. Its seeds are comprised of oil (45–60%), proteins (18-25%), carbohydrate (3-25%), and rich with essential vitamins and mineral-nutrients (Anilakumar et al., 2010; Teboul et al., 2020). The seeds are being used for an array of products in the food (e.g., high-quality oil, tahini paste, and cooking and backing) and pharmaceutical industries (Mushtaq, 2020). At present, sesame is cultivated mainly in developing countries, with annual seed production of 6.7 million tons (http://www.fao.org/faostat/en/#data/QC). The global shift toward healthier and more nutritional plant-based food products lead to significantly increased demand for sesame seeds and derivative products. However, despite its economical and agricultural importance, sesame is considered an ‘orphan crop-plant’ and has been subjected to limited agronomical and scientific research.

Sesame is a short-day erect plant with an indeterminate florescence and simple or branching rigid stem. Its growth period ranges usually from 12-18 weeks, with flowering (i.e., the transition from vegetative to reproductive phase) begins about 30-40 (early) to 70-80 (late) days after sowing (Langham, 2007). This variation in flowering time could affect the crop adaption to specific agro-system conditions. As the blooming period continues until plant maturation, the flowering date plays a crucial role in both plant architecture and yield components. Sesame yield components include the number of plants per unit area, number of branches per plant, number of capsules per leaf axil, seeds per capsule, and seed weight (York & Garden, 2017). Complex tradeoffs between yield component traits have been shown to significantly affect the final seed yield (Gadri et al., 2020). Branching habit and number of capsules per leaf axil were shown to support higher seed yield (Mei et al., 2017), whereas dwarf mutants (i.e., small plant height) negatively affect seed weight (Miao et al., 2020).

The advent of next-generation sequencing and genotyping by sequencing (GBS) technologies has provided a means for examining genetic diversity and population structure of crop-plants, which can facilitate the genetic dissection of agronomic traits and integrate them in breeding programs. Genome-wide association studies (GWAS) are a promising approach that connects phenotypic variation and genomic data (i.e., genetic markers) to detect genomic regions underlying complex traits (Tibbs Cortes et al., 2021). GWAS were applied successfully for various crop-plants, such as bread wheat (*Triticum aestivum* L.; Guo et al., 2017), maize (*Zea mays* L.; Li et al., 2012), rice (*Oryza sativa* L.; Zhao et al., 2011), and soybean (*Glycine max* L.; Sonah, 2015). In sesame, GWAS were used for the identification of genomic regions associated with response to biotic (Asekova et al., 2021) and abiotic (Li et al., 2018; Dossa et al., 2019) stresses, as well as yield-related traits (Zhou et al., 2018). The relatively small genome size (~375 Mbps), the recent development of genomic resources and rich genetic diversity make sesame an ideal model crop for genetic investigation (Dossa et al., 2017).

Here we harness the natural variation among geographically distributed sesame germplasm to underpin the genetic architecture of morpho-agronomic and yield-related traits. Our working hypothesis was that as a consequence of minimal artificial selection processes (associated with modern breeding), sesame preserved rich genetic and phenotypic diversity that will enable detection of novel genomic regions conferring agronomical important traits. The aims of the current study were to (***i***) characterize the genetic diversity in the newly established sesame collection, (***ii***) detect genomic regions contributing to the phenotypic performance, and (***iii***) infer the genetic associations between traits. Our findings shed new light on the interaction between flowering date, morpho-physiological traits, and yield components in sesame.

## 2. MATERIALS AND METHODS

### 2.1 Plant material and experimental design

A panel of 184 sesame genotypes from the Hebrew University of Jerusalem sesame collection was assembled (SCHUJI panel, hereafter) according to their geographical origins to capture the whole sesame genepool genetic diversity (Supplemental Table S1). The plants were grown over two growing seasons (2018 and 2020) at the experimental farm of Hebrew University of Jerusalem in Rehovot, Israel (34°47′N, 31°54′E; 54 m above sea level). The soil at this location is brown-red degrading sandy loam (Rhodoxeralf) composed of 76% sand, 8% silt, and 16% clay. In the 2018 growing season, a complete random factorial (genotypes) block design with seven replicates was employed. Each block consisted of 184 plots sown as single row, with six plants, 15cm apart (1-meter-long plot). The two plants at the edges of each plot served as borders. The remaining four plants were used for phenotypic characterization and at the end of the experiment, they were harvested to estimate yield components. In the 2020 growing season, the same experimental design was employed with five replicates per genotype. The plot size was 2.6 m × 0.8 m (15 cm between plants) with three rows per plot. Five representing plants from the middle row were used for phenotypic evaluation at maturity and harvested to obtain yield components. In both seasons, the field was treated with fungicides and pesticides to avoid the development of fungal pathogens or insect pests and was weeded manually once a week.

### 2.2 Phenotypic measurements

Phenotypes were recorded during the whole sesame growing season for each plot. *Flowering date* (FD) was evaluated visually when 50% of the plants in each plot had at least one open flower. *Height to the first capsule* (HTFC) and *plant height* (PH) were measured at maturity from the soil surface to the first capsule and the plant tip, respectively. The *reproductive zone of the main stem* (RZ) was calculated as the delta between PH and HTFC, and the *reproductive index* (RI) was calculated as the ratio between RZ and PH (RZ/PH). Before harvest, the *number of branches per plant* (NBPP) were counted as an average of all individual plant in a plot. At physiological maturity, three plants (2018) and five plants (2020) from each plot were harvested, and sun-dried. The samples were threshed using the laboratory threshing machine (LD 350, WinterSteiger, Reid, Austria). Seeds were counted using the seeds counting machine (Data Count S25, Data Technologies) and weighted in analytical lab weight to obtain *seed number per plant* (SNPP), *seed yield per plant* (SYPP), and *thousand-seed weight* (TSW) for each plot.

### 2.3 Statistical analysis of phenotypic data

The JMP ver.15 pro statistical package (SAS Institute, Cary, NC, USA) and R (R Core Team, 2020) was used for all statistical analyses with a significant threshold of 5%. First, we calculated the best linear unbiased estimate (BLUE) for every trait for each genotype per year and for both years using the lme4 R package (Bates et al., 2015). The mixed linear model for BLUE per year was fitted according to the formula:

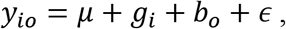

where *y_io_* is the phenotypic observation for the *i*th genotype in the *o*th block, *μ* is the intercept, *g_i_* is the genotype fixed effect, *b_o_* is the block random effect, and *ϵ* is the model residuals.

The BLUE for the combined data from the two years was calculated according to the formula:

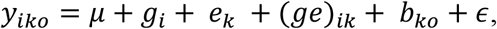

where *y_iko_* is the phenotypic observation for the *i*th genotype in the *k*th year and the *o*th block, *μ* is the intercept, *g_i_* is the genotype fixed effect, *e_k_* is the random effect of year, (*ge*)_*ik*_ is the random effect of genotype-by-year interaction, *b_ko_* is the random effect of block nested within year, and *ϵ* is the model residual. For the calculation of heritability, we fitted the same mixed model as above, with the expectation of genotype, that considered as random effect. A broad-sense heritability was calculated on the entry-mean basis according to Schmidt et al. (2019) using the estimated variance components:

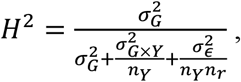

where 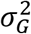, 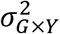 and 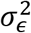 are the genetic, genotype-by-year interaction, and residual variances, respectively, and *n_Y_* is the number of years (2) and *n_r_* is the average number of replicates across years (6). The significance of variance components was evaluated by the likelihood ratio test using the lmerTest R package (Kuznetsova et al., 2017).

We performed GWAS for each data set (referred to as 2018 data, 2020 data, and combined data, hereafter). The combined data set was used to infer phenotypic and genomic correlations and to conduct principal components analysis (PCA), k-means clustering, and genomic heritability estimation. K-means clustering analysis was applied using the factoextra R package (Kassambara and Mundt, 2020) on the centered and scaled values of each trait. Both PCA and clusters plot was drawn in the JMP (ver.15pro) statistical package. Density plots and a correlation matrix were constructed using the ggplot2 (Wickham, 2016) and the corrplot (Wei and Simko, 2017) R packages.

### 2.4 Genotyping and preparation of marker data set

Genomic DNA was extracted from young leaf tissues with a modified CTAB method (Doyle, 1987). We generated GBS data using the procedure described in Elshire et al. (2011), with minor changes: 100 ng of genomic DNA and 3.6 ng of total adapters were used. Genomic DNA was restricted with ApeKI enzyme and the library was amplified with 18 PCR cycles. We used the Zhongzhi No. 13 (https://www.ncbi.nlm.nih.gov/assembly/GCF_000512975.1) reference genome to perform reference-based SNP calls based on the STACKS 2.3 pipeline (http://catchenlab.life.illinois.edu/stacks). These analyses yielded 90,542 single nucleotide polymorphisms markers (SNPs) in total including unknown scaffolds and chloroplast genome. Markers on unknown scaffolds and chloroplast genome were removed and all the markers were filtered to depth quality of 3 using TASSEL ver. 5.0 (Bradbury et al., 2007). Imputation of missing genotypes was performed by BEAGLE ver. 5 (Browning & Browning, 2007). We also removed markers that were tightly linked (r^2^=0.99) using PLINK (Purcell et al., 2007). Polymorphic sites with <5% minor allele frequency and > 20% heterozygosity were filtered out by PLINK and TASSEL, respectively. The remaining 20,294 SNPs were used for further analysis.

### 2.5 Population structure, kinship, and linkage disequilibrium

We used PCA and a centered identity-by-state matrix (G) constructed from TASSEL (Bradbury et al., 2007) to infer population structure in the sesame panel. The ADMIXTURE software (Alexander et al., 2009) was used to estimate a Q-matrix, which is the ancestry among the accessions. To select the number of subpopulations (K), we ran the software from K=1 to 10 with five-fold cross-validation. This analysis outputs the cross-validation error (%) for a given K. The number of subpopulations was determined as K that produced the lowest cross-validation error. The lowest value of cross-validation error was achieved with K=7 (Supplemental Table S2), and the Q-matrix was constructed with 1,000 bootstrapping. The results from the PCA analysis and ADMIXTURE software outputs were plotted using the ggplot2 R package. The fixation index (F_st_) among the seven subpopulations was calculated with VCFtools (Danecek et al., 2011) according to Weir & Cockerham (1984). Genome-wide linkage disequilibrium (LD) was obtained through pairwise correlations between markers with a sliding window of 10 markers using PLINK (Purcell et al., 2007). The correlation between markers (r^2^) was plotted against their physical positions (i.e., base pairs) and the extent of LD pattern and decay was obtained by fitting a non-linear model according to Hill & Weir (1988) as described in Marroni et al. (2011). We used a value (in base pairs) in which LD halves from its initial value for defining haplotypes and mining for candidate genes (CG) around significant markers.

### 2.6 Genomic heritability and genomic correlations

Genomic heritability estimates and genomic correlations were inferred using the BGLR R package (Pérez & De Los Campos, 2014). To obtain genomic heritability for each trait, we fitted a Bayesian univariate genomic best linear unbiased prediction using the equation:

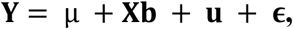

where **Y** is the vector of single-trait BLUE phenotypes, μ is the intercept, **X** is a design matrix for fixed effects, **b** is the vector of fixed effects containing three PCs to account for population structure, **u** is the vector of random effects, and **ϵ** is a vector of the model residuals. The following distributions were assumed for random effects:

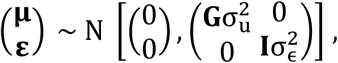

where **G** represents the first genomic relationship matrix of VanRaden (2008), **I** is the identity matrix, and 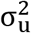 is the additive genomic variance explained by genetic markers, and 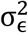 is the model residuals. Genomic heritability was calculated as: 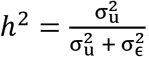.

Genomic correlations were estimated using the multivariate version of the model described above, where **Y** is the vector of multi-trait phenotypes. The following distributions were assumed for random effects:

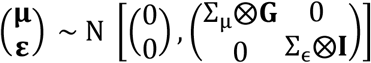

where Σu and Σe refer to the genetic and residual variance-covariance matrices, respectively, and ⊗ is the Kronecker product.

Genomic correlations between traits were derived from the genomic variance-covariance matrix as:

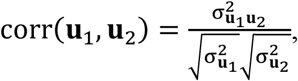

where **u**_1_ and **u**_2_ are the breeding values of traits **Y**_1_ and **Y**_2_, 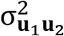 is the additive genomic covariance between **u**_1_ and **u**_2_, and 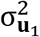 and 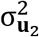 are additive genomic variances for **Y**_1_ and **Y**_2_, respectively.

### 2.7 Association mapping

To identify genomic regions associated with the traits of interest, we used a mixed linear model of Henderson (1975) coupled with the first three PCs and G matrices to account for population structure and relatedness among individuals, respectively, using the rrBLUP R package (Endelman, 2011). We fit an additive single-marker GWAS model as the following:

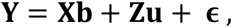

where **Y** is the vector of phenotypes, **b** is the vector of fixed effects including an SNP and 3 PCs, **u** is a vector of random additive genetic effects with mean zero and variance-covariance 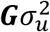, **X** and **Z** are the known incidence matrices, and **ϵ** is the vector of residuals (Yu et al., 2006). The *P*-value threshold (1.551509 × 10^-5^) was determined by calculating the number of effective independent tests (Meff) as described in Li & Ji (2005) with the following formula: *P* = 1 - (1-0.05)^(1/Meff), where *P* is the genome-wide *P*-value threshold and 0.05 is the desired level of significance, and Meff = 3,306.

### 2.8 Haplotypes estimation and candidate gene analysis

Haplotypes encompassing associated markers that corresponded to the LD-decay pattern in the SCHUJI panel was constructed in TASSEL version 5.0 (Bradbury et al., 2007). We included only haplotypes with a frequency greater than 5% and only genotypes that were homozygous in all the genetic markers within a haplotype. The effects of haplotypes on the phenotypic variation were estimated using analysis of variance (JMP ver. 15) at a significant level of 5%. Phenotypic response was the BLUE of 2018 and 2020 growing seasons. A haplotype plot was produced using the Rainclouds R package (Allen et al., 2021). We used LD-decay for mining CGs around significant SNPs and analyzed them according to the Zhongzhi No. 13 reference genome (https://www.ncbi.nlm.nih.gov/assembly/GCF_000512975.1).

## 3. RESULTS

### 3.1 High phenotypic diversity among the sesame panel

To test the level of phenotypic diversity among the newly established SCHUJI panel, we characterized the sesame under Mediterranean basin conditions over two seasons. In general, the SCHUJI panel exhibited rich variation for phenological, plant architecture, and yield components (Fig. 1; Supplemental Table S3; Fig. S1). Flowering date (FD) showed a similar pattern across years and spread along most of the growing season, ranging between 39.86 to 76 and 39.2 to 70 days after sowing (DAS) for 2018 and 2020, respectively. In both years (2018 and 2020), most genotypes (135 and 146) flowered before 55 DAS (Fig. 1A). Plant architecture traits showed a wider range in the 2020 than 2018 seasons for HTFC (29-140 *vs*. 36-198 cm; Fig. 1B), PH (80-177 *vs*. 103-251 cm; Fig. 1C), and RZ (20 to 104 *vs*. 28 to 141cm; Fig. S1). In contrast, RI exhibited a similar range in both seasons (0.13-0.73 vs. 0.12-0.76; Fig. 1D). Most yield components showed a similar pattern across years (SYPP and TSW) except SNPP which exhibited a much wider variation in 2020 (21-7719 *vs*. 508-13591) (Fig. 1F-H).

**Figure 1.**
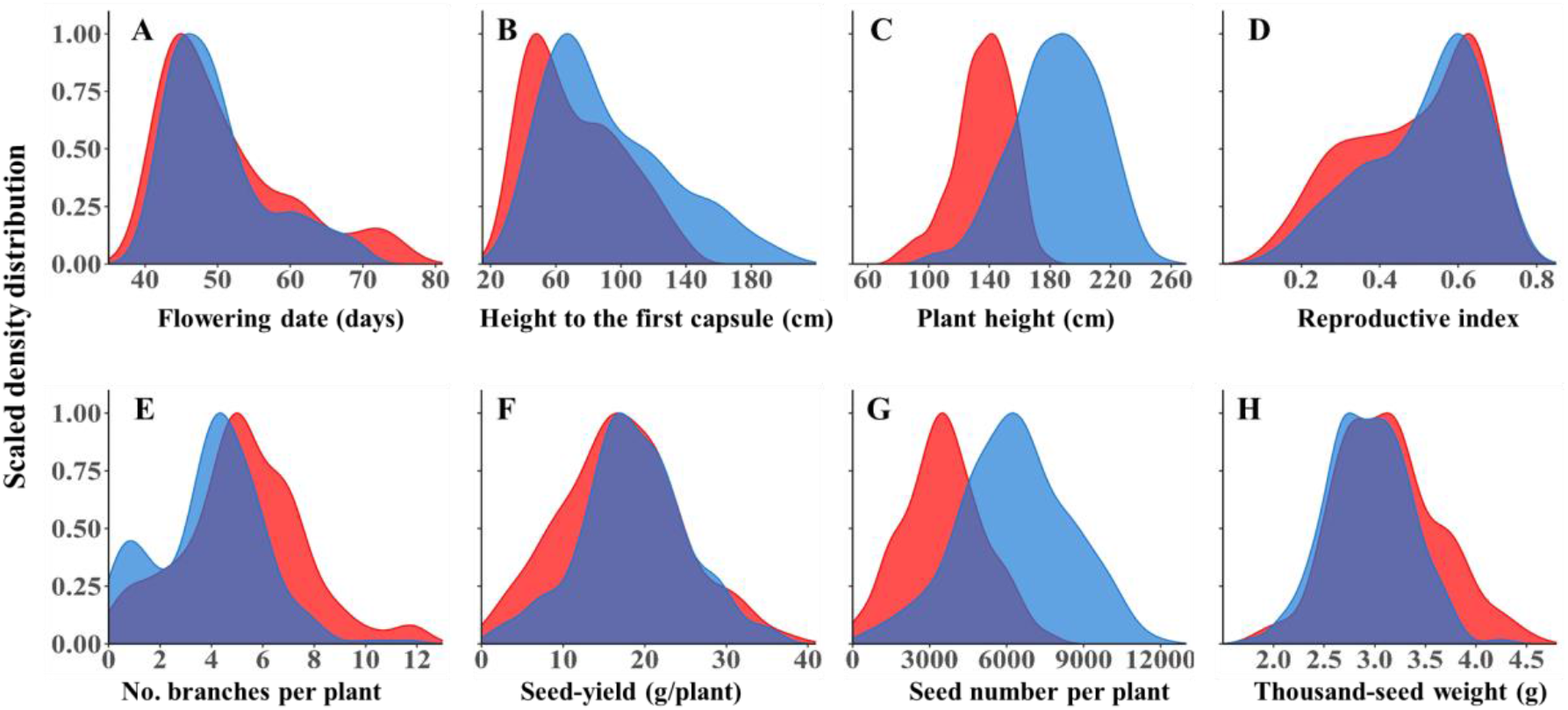
Density distribution of *phenological traits*: (**A**) Flowering date. *Plant architecture traits:* (**B**) height to the first capsule, (**C**) plant height, (**D**) reproductive index, and (**E**) number of branches per plant. *Yield components:* (**F**) seed yield per plant, (**G**) number of seeds per plant, and (**H**) thousand-seed weight, under two years: 2018 (red) and 2020 (blue).

To test the effect of genotype (G), year (Y), and genotype-by-year interaction (G×Y), we estimated the variance component of each parameter for all the traits. In general, genotype and year had significant effects for most of the traits, with a significant G×Y interaction. The estimates of broad-sense heritability ranged from moderate (0.66) for SNPP to high (0.97) for FD (Table 1).

**Table 1.** Variance component estimates for the random effects. σ^2^_G_ is the genetic variance, σ^2^_Y_ is the year variance, σ^2^_G×Y_ is the genotype-by-year variance, 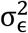 is the error variance, and *H*^2^ is the broad-sense heritability. Flowering date (FD), height to the first capsule (HTFC), plant height (PH), number of branches per plant (NBPP), reproductive index (RI), seed yield per plant (SYPP), number of seeds per plant (SNPP), and thousand-seed weight (TSW).

| Trait | Variance component | | | | $H^2$ |
| --- | --- | --- | --- | --- | --- |
| | $\sigma^2_G$ | $\sigma^2_Y$ | $\sigma^2_{G \times Y}$ | $\sigma^2_\epsilon$ | |
| **FD** | 61.48*** | 0.39<sup>n.s.</sup> | 2.77*** | 8.84 | 0.97 |
| **HTFC** | 1069 *** | 214.6*** | 71.49*** | 125.95 | 0.96 |
| **PH** | 441.81*** | 1174.8*** | 44.18*** | 289.12 | 0.89 |
| **RZ** | 356.97*** | 394.51*** | 31.97*** | 204.56 | 0.91 |
| **RI** | 0.02*** | 0.0003* | 0.0009*** | 0.003 | 0.97 |
| **NBPP** | 4.04*** | 0.72** | 0.64*** | 2.83 | 0.88 |
| **SYPP** | 28.36*** | 0.72<sup>n.s.</sup> | 10.21*** | 54.66 | 0.74 |
| **SNPP** | 1723700.7*** | 3696689.4* | 854375.3*** | 5260739 | 0.66 |
| **TSW** | 0.17*** | 0.02*** | 0.03*** | 0.05 | 0.88 |

n.s., \* and \*\*\* indicate non significant, significant differences at $P \leq 0.05$ and 0.001, respectively, as determined by the likelihood ratio test.

To examine the relationship between vegetative-related traits (late FD, PH, and NBPP) and yield-related traits (RI and yield components), we applied PCA to BLUEs from the combined data. Based on this analysis, three PCs (eigenvalues > 1) accounted for 75.1% of the total phenotypic variance among the genotypes (Fig. 2A). PC1 explained 54.3% of the variation, loaded positively to FD, HTFC, PH, and NBPP, and negatively loaded to RI, SNPP, SYPP, and TSW. PC2 explained 20.8% of the variation and loaded positively to HTFC, PH, NBPP, SYPP, SNPP, and TSW and negatively loaded to FD and RI. SYPP was positively correlated with SNPP (r=0.86) and TSW (r=0.37), however, no significant relationship was observed between these two traits (r=-0.04; Supplemental Table S4). Overall, FD and PH were positively correlated (r=0.7) and both were negatively correlated with RI (r=-0.89 and r=-0.68, respectively; Fig. 2A; Supplemental Table S4).

**Figure 2.**
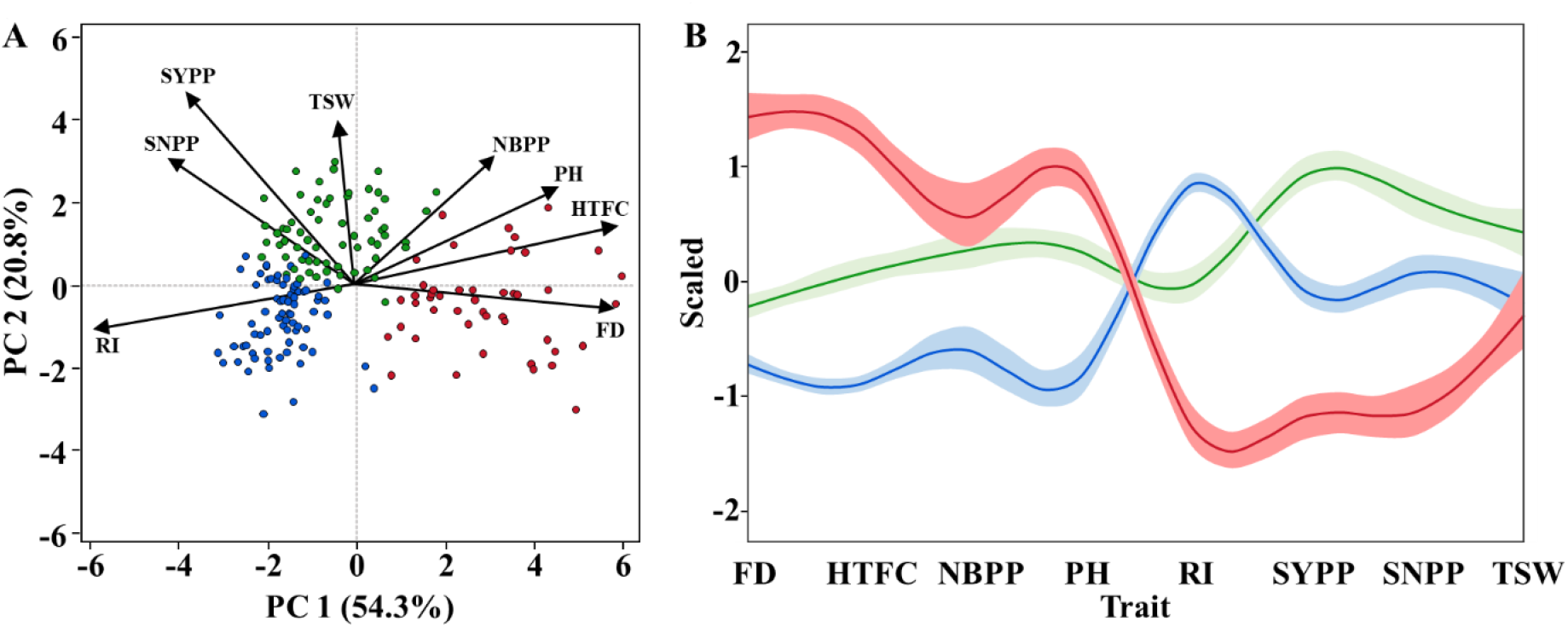
(**A**) Principal component (PC) analysis of phenotypic traits of the combined data (2018 and 2020). Each dot represents one genotype. (**B**) K-means clustering analysis of the primary traits. Y-axis is the centered and scaled values for each trait and the lines are the cluster means with their confidence intervals. Clusters 1 (Green), 2, (Blue), and 3 (red) were estimated using the K-means clustering analysis. The traits included Flowering date (FD), height to the first capsule (HTFC), plant height (PH), number of branches per plant (NBPP), reproductive index (RI), seed yield per plant (SYPP), number of seeds per plant (SNPP), and thousand-seed weight (TSW).

### 3.2 Flowering date affects the final seed-yield via morphological modifications

To further dissect the relationship between flowering date, plant architecture, and yield components, we applied k-means clustering analysis using the combined data. This analysis partitioned the panel into three clusters: early flowering (average 44.84 DAS), mid flowering (48.76 DAS), and late flowering (61.65 DAS) (Fig. 2B; Supplemental Table S5). Comparison between clusters 2 and 1 (early and mid-flowering) shows that they are different in terms of morphological and yield components. Although cluster 2 genotypes have larger RI (0.63 *vs*. 0.5), genotypes from cluster 1 had greater PH and NBPP (142.23 *vs*. 166.69 cm and 3.29 *vs*. 5.12 branches per plant, respectively). These alternations in the plant architecture traits affect the outcome, as we observed higher SYPP, SNPP, and TSW for cluster 1 genotypes (Fig. 2B; Supplemental Table S5). When we compared clusters 1 and 2 to cluster 3, we observed a major difference in FD (61.65 days) that led to a long vegetative phase and higher PH (181.77 cm) and NBPP (5.67), but these cluster 3 genotypes had lower RI (0.31) and lower SNPP, SYPP, and TSW performance.

### 3.3 Allelic diversity and population structure of the sesame panel

To examine the genetic diversity at the genomic level, we used 20,294 SNP markers, spread along the sesame genome. Principal component analysis on the SNP data explained 30.8% of the genetic variation between genotypes. The PCA did not show any separation between genotypes relative to their geographical origins (Fig. 3A). ADMIXTURE analysis partitioned the panel into four major (K1, K5, K6, and K7) and three minor (K2, K3, and K4) sub-populations (Fig. 3B). F_st_ values of these seven sub-populations ranged from 0.08 to 0.32, which showed a weak to moderate differentiation among the subpopulations (Supplemental Table S6). Genome-wide LD analysis showed that LD decayed rapidly to half of its initial value (0.22) at 58,774 base pairs (Supplemental Fig. S2).

**Figure 3.**
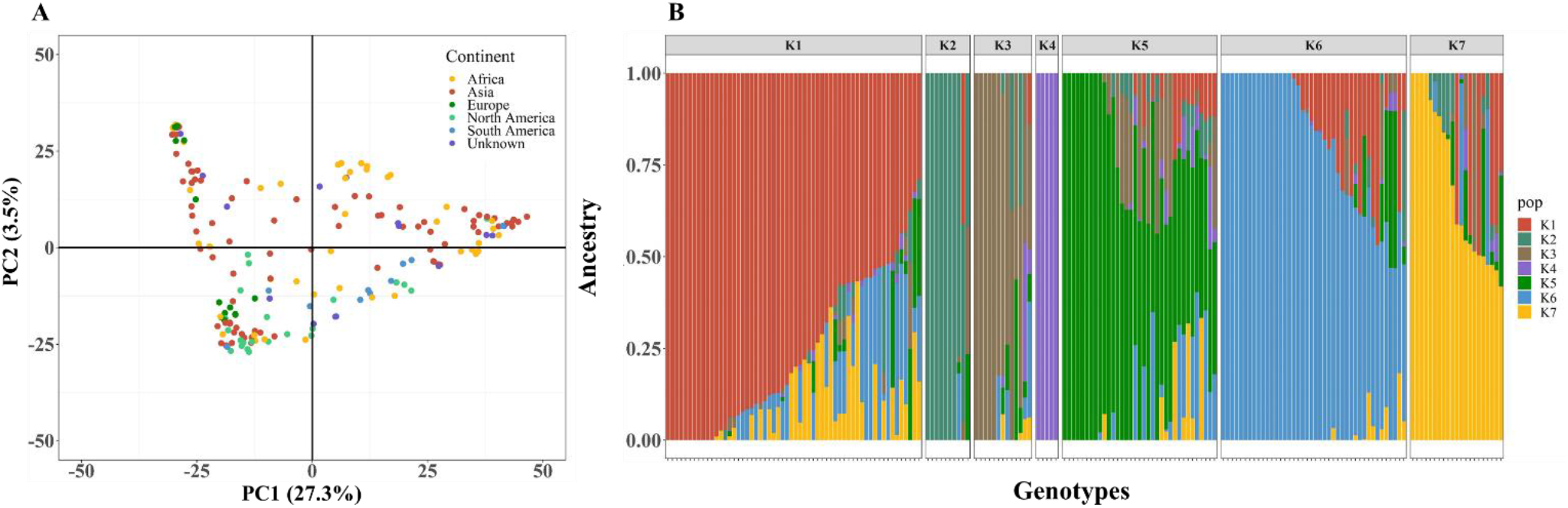
Population structure of the SCHUJI panel. (**A**) Principal component analysis and (**B**) ADMIXTURE when K=7. Every single dot or line represents an individual genotype.

### 3.4 Genomic heritability and genomic correlations

To elucidate the trait similarity at the genomic level, we computed genomic heritability for each trait and genomic correlations between the measured traits. Estimates of genomic heritability ranged from 0.37 (PH) to 0.58 (SYPP) presenting moderate values for all the traits (Supplemental Table S7). Figure 4 presents phenotypic (upper triangular elements) and genomic (lower triangular elements) correlations between the traits. Overall, the phenotypic and genomic correlations showed similar patterns (Fig. 4; supplemental Tables S4, S8). FD was positively correlated with HTFC and PH (0.88 and 0.58, respectively), while negatively correlated with morphological yield-related traits, such as RZ and RI. HTFC was found to be positively correlated with PH (0.71) while negativity correlated with RZ (−0.75), RI (−0.96), SYPP (−0.56), and SNPP (−0.6). SYPP was positively correlated with SNPP (0.93) and TSW (0.44), whereas SNPP and TSW seemed to be less related (0.17).

**Figure 4.**
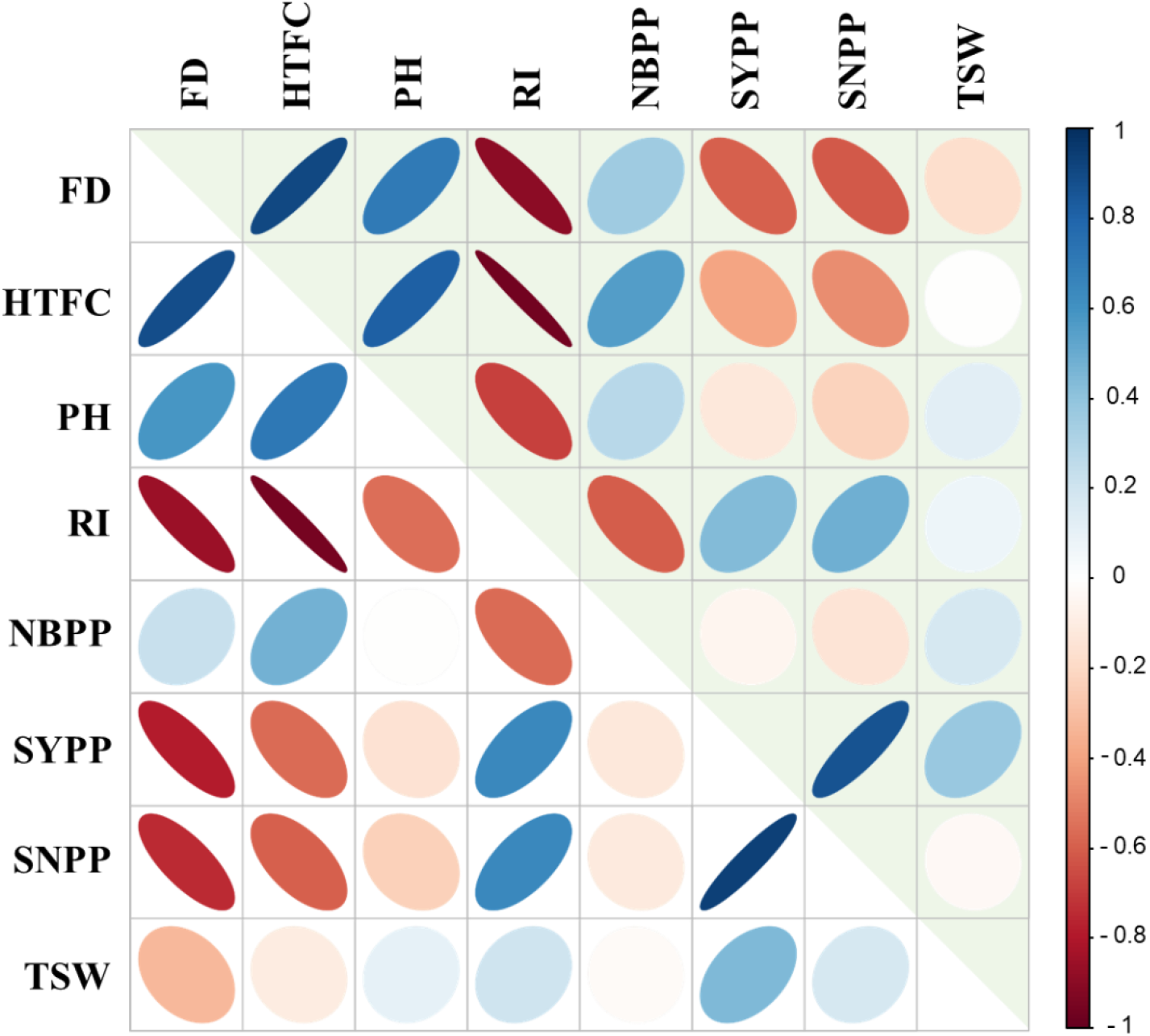
Heatmap of phenotypic (upper triangular elements) and genomic (lower triangular elements) correlation matrix between the primary traits: Flowering date (FD), height to the first capsule (HTFC), plant height (PH), reproductive index (RI), number of branches per plant (NBPP),), seed yield per plant (SYPP), number of seeds per plant (SNPP), and thousand-seed weight (TSW). Colors indicate the level of correlations (r) from positive (blue) to negative (red).

### 3.5 Association mapping of agronomic traits

We performed a single-marker regression GWAS for all the traits and detected 11, 19, and 20 SNPs that are associated with trait mean differences for 2018, 2020, and combined data, respectively (Supplemental Table S9). For FD, we identified two major genomic regions on linkage group (LG) 2 and 11 (4, 4, and 5 SNPs for 2018, 2020 and combined data, respectively; Fig. 5A). In total, 9 SNPs were found significantly associated with the plant architecture traits, including 4 for HTFC (Fig. 5B), 3 for RI (Fig. 5C), and 2 for RZ (Supplemental Fig. S3A). For PH and NBPP, there was no clear peak (Supplemental Fig. S3B-C).

**Figure 5.**
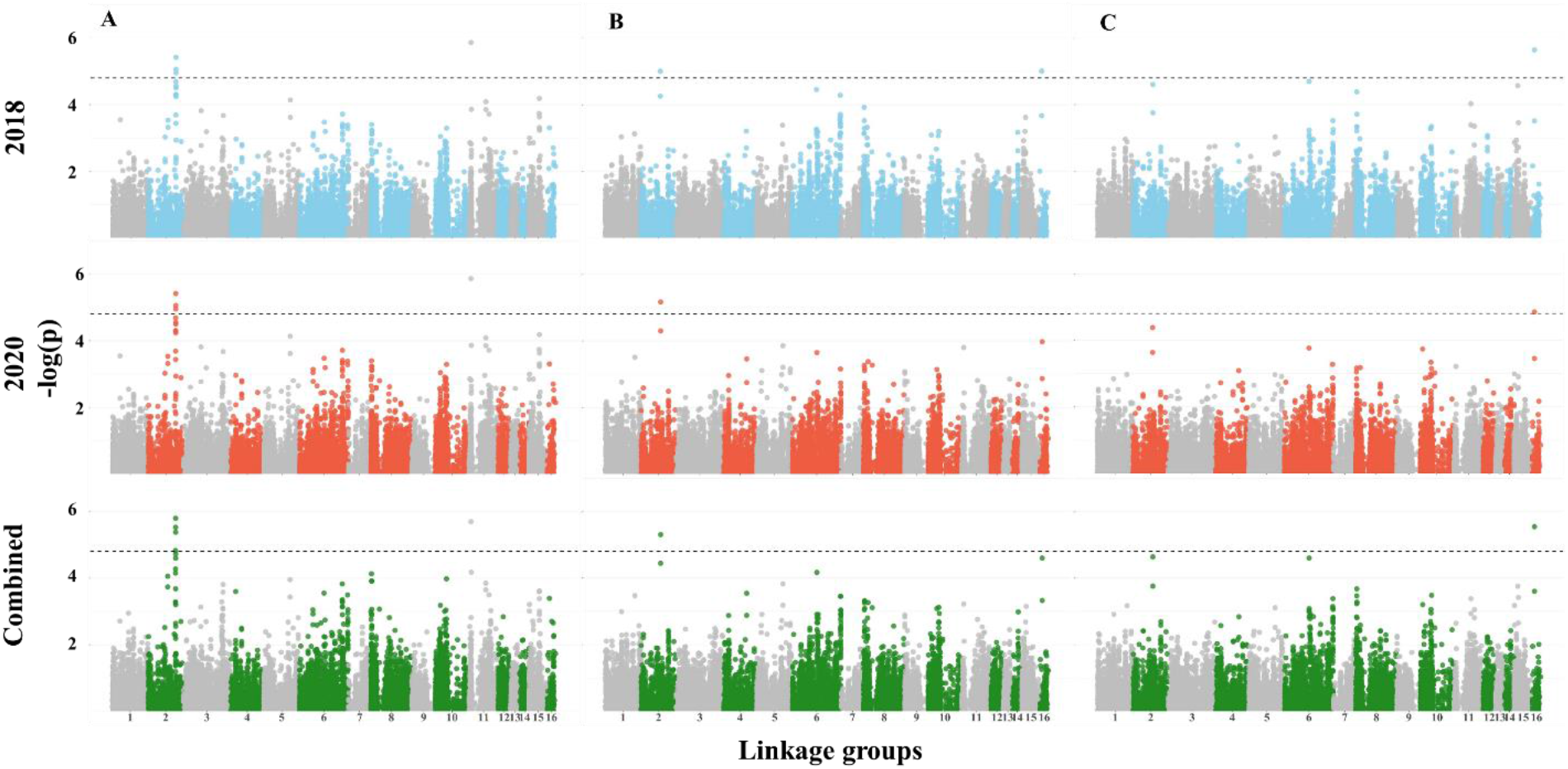
Manhattan plots for (**A)** flowering date, (**B**) height to the first capsule, and (**C**) reproductive index in 2018, 2020, and combined data. The dashed line represents the genome-wide significance threshold.

For SYPP and SNPP, we detected one major genomic region on LG2 (except for SNPP in 2018) (Fig. 6A-B). SYPP was associated with 20 SNPs while SNPP was associated with 7 SNPs (Supplemental Table S9). For TSW, we found one SNPs on LG1 that was slightly below the significant threshold (-log_10_(*p*) = 4.77), only in the 2020 season (Fig. 6C, and Supplemental Table S9). We compared the mapping results to find SNPs that overlap across traits. HTFC had one genomic region overlapping with RI on LG16 (Supplemental Table S9). FD had a major genomic region overlapping with SYPP and SNPP (Figs. 5-6; Supplemental Table S9).

**Figure 6.**
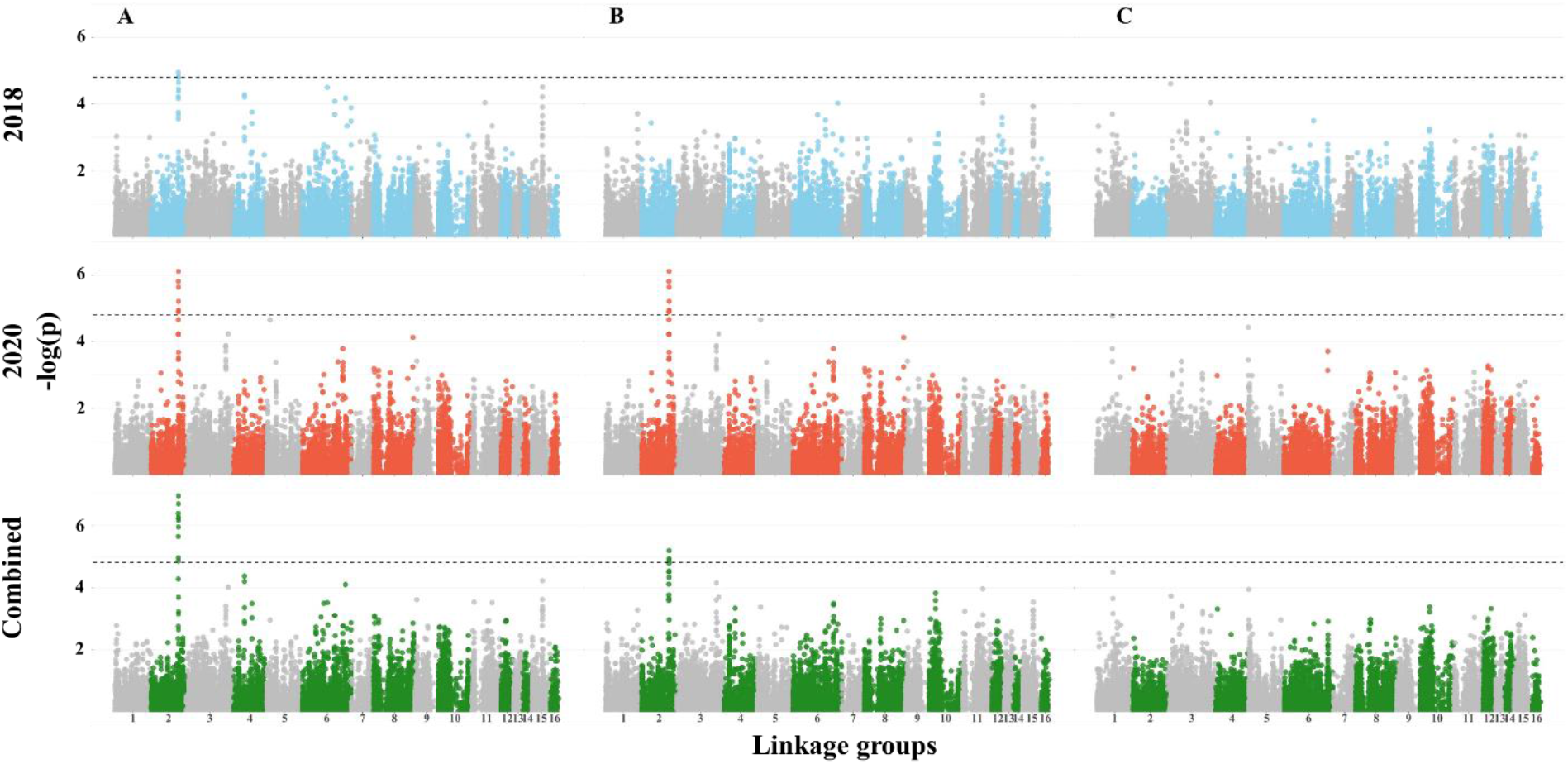
Manhattan plots for yield components traits: (**A**) seed yield per plant, (**B**) seed number per plant, and (**C**) thousand-seed weight. The dashed line represents the genome-wide significance threshold.

### 3.7 Flowering date promotes yield stability in sesame

A significant genomic region in the length of 96,491 base pairs that contained 10 SNPs on LG2 was found to be associated with FD, SNPP, and SYPP (Supplemental Table S9). In total, 4 SNPs inside this region overlapped with FD and SYPP and were within the range of LD-decay (Supplemental Fig. S2) in our panel. To explore their influence on the phenotypic variation of these two traits across the two growing seasons, we defined haplotypes for these 4 overlapping SNPs. Two possible haplotypes were found. The first haplotype (Hap1) was more frequent (*n*=140) than the second haplotype (Hap2, *n*=30) (Fig. 7). Hap1 was found associated with earliness while genotypes that included Hap2 exhibited late flowering under the Mediterranean climate (*P*<0.0001, Fig. 7A). Moreover, these two haplotypes also differed in yield performance, with Hap1 promoting higher seed yield (Fig. 7B). To test the phenotypic stability of the haplotypes across years, we perform an analysis of variance for haplotypes, year, and their interaction using FD and SYPP from BLUE per year analysis. While the haplotypes had a significant effect on the traits (*P*<0.0001 for FD and SYPP), we did not observe any significant interaction between haplotypes and year (*P*=0.087 for FD and *P*=0.82 for SYPP). These results may indicate that this genomic region promotes yield stability via modifications in FD. Moreover, we obtained differences in phenotypic responses within haplotypes (per genotype) in FD and SYPP across years. For FD, we found that genotypes in both haplotypes had a similar pattern (*P*=0.5 and *P*=0.12, for year effect) as expressed in parallel trend lines between genotypes across years (Fig. 7A). For SYPP, the differences in mean values across years were on the edge of statistical significance (*P*=0.055 and *P*=0.32 for Hap1 and Hap2, respectively), but the genotypes within each haplotype had different values across years as expressed with the crossing lines that connected the same genotypes between the two growing seasons (Fig. 7B).

**Figure 7.**
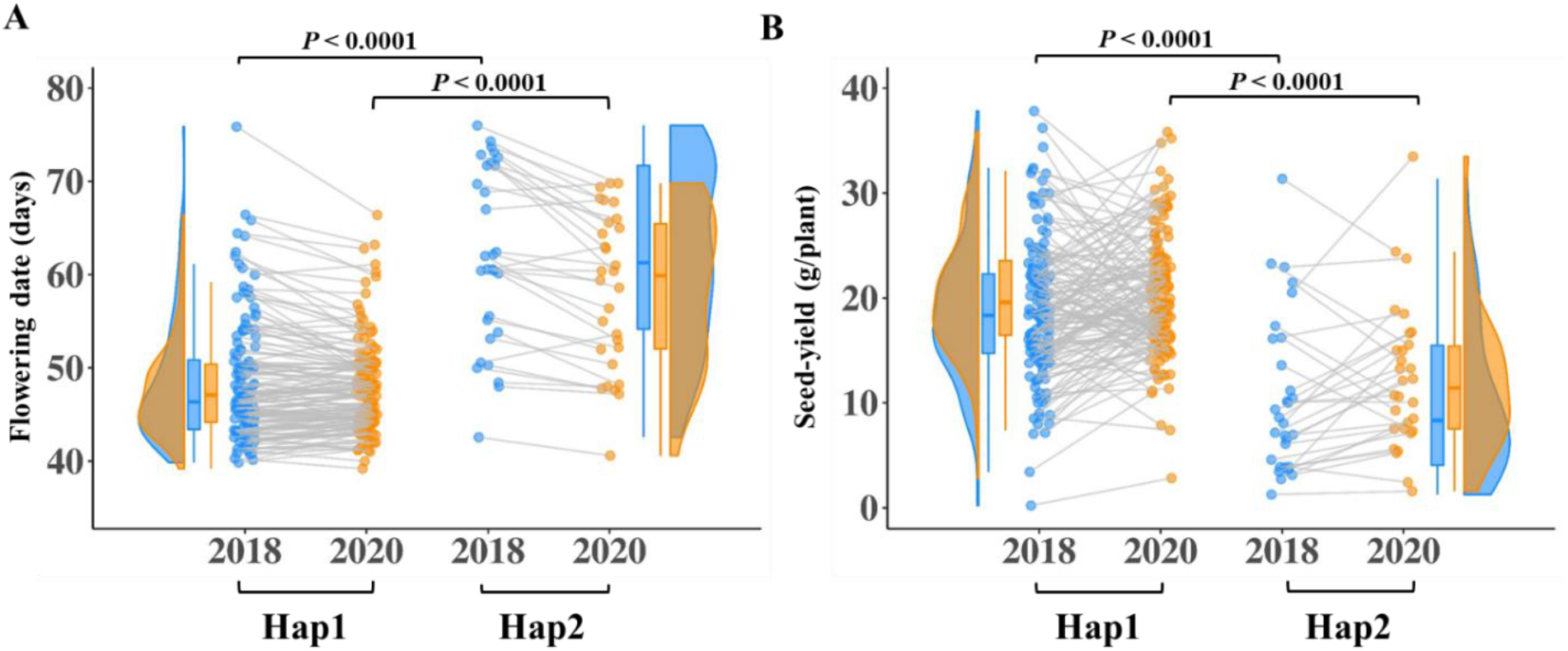
Haplotype analysis of the major genomic region on LG2 and their effects on (**A**) flowering date and (**B**) seed yield across two years. Every dot represents a genotype. Blue color represents the 2018 growing season, orange represents the 2020 growing season, and grey lines connect the same genotypes in the two growing seasons.

To find the genetic basis underlying the biological influence on the traits, we searched for candidate genes (CGs) within this LG2 hotspot genomic region. Overall, we found 20 CGs: 7 CGs were located inside of the genomic region, while 13 CGs were consisted of upstream (8) and downstream (5) of the genomic region. (Supplemental Table S10). Although four of the CGs encode uncharacterized proteins, most of the CGs were involved in signaling pathways and regulation and four of them (LOC105156148, LOC105156152, LOC105156159, and LOC105156313) were associated with controlling flowering time, floral development, and productivity in *Arabidopsis* and rice (Ito et al., 2007; Wu et al., 2012; Miao et al., 2018; Teng et al., 2019).

## 4. DISCUSSION

Understanding the genetic basis of agronomic traits is important for research and breeding efforts. Here, we explored the genetic and phenotypic variation in a newly established sesame panel (SCHUJI) aiming at dissecting the genomic architecture of morpho-agronomic traits. We hypothesized that as ‘orphan crop-plant’ sesame has not been subjected to modern selection, the geographically distributed panel preserved an ample allelic repertoire for agronomically important traits. Evaluation of the level and extent of genetic diversity among the sesame panel surprisingly showed that there is no clear geographical separation between genotypes (Fig. 3A). Similarly, previous genetic characterization of other sesame panels showed no clustering associated with continents and suggested sub-geographical regions or latitudes as key factors (Wei et al., 2015; Cai et al., 2016; Dossa et al., 2016). Moreover, the ADMIXTURE analysis resulted in seven sub-populations (K=7) with a low to moderate diffrrences between them (Fig. 3B; Supplemental Table S6). Thus, the lack of geographic signature in our, as well as other genetic panels, may be a consequence of recent genetic materials exchange (Basak et al., 2019) or unavailable information at the collection sites. As an example, even the smallest sub-populations (2, 3, and 4) that were more genetically uniform showed the various origin of genotypes (i.e., sub-population 4 included genotypes from Thailand, Israel, or unknown) (Fig. 3B).

### 4.1 Wide phenotypic diversity among the sesame panel highlights the tradeoff between flowering date and productivity traits

High phenotypic diversity was found for all morpho-agronomical traits across the two years, with 2020 expressing higher values (Fig. 1). This observation agrees with previous studies using other sesame panels (Furat & Uzun, 2010; Zhou et al., 2018). In general, FD was found to be negatively correlated with yield components, such as RI, SYPP, and SNPP. On the other hand, late FD was associated with high NBPP, HTFC, and PH (Figs. 2, 4; Supplemental Table S4). Additional support for the key effect of FD on productivity comes from the k-means clustering analysis. This analysis clustered the panel into three groups (Fig. 2B), which were also associated with the genetic structure of the panel (Supplemental Fig. S4). Comparison between the mid-FD group (green) and the early FD (blue) showed that NBPP and PH have a positive effect on productivity traits, which is in line with previous reports in sesame (Baydar, 2005). Gadri et al. (2020) showed in sesame that increasing the source size (i.e., vegetative organs) can support a higher seed set and filling. Thus, the higher yield potential of cluster 1 can be a consequence of greater vegetative biomass accumulation following more branches. Interestingly, the late-flowering group (red) had a similar number of branches as the intermediate group (green); however, most of them were not fertile due to the late-flowering phenotype. RI is a key trait for yield potential assessment and represents the ratio between RZ and PH. As expected, RI and FD are negatively correlated (Fig. 2). However, it is worth noted that a high value of RI can be a consequence of either high PH with low HTFC or small plants. Therefore, to promote higher yield potential, it is important to combine high RI with high RZ values.

Langham (2007) reported that photoperiod responsiveness plays a key role in sesame flowering date and vegetative biomass accumulation. Our results show that while most genotypes that belong to the late FD and low RI group clustered together, they differ in their geographical origins (Fig. 3A; Supplemental Fig. S4), which may suggest the involvement of other genetic and/or enviomental factors.

### 4.2 Genetic architecture of agronomical traits reveals hotspot of overlapping genomic regions

To detect the genetic basis underlying observed phenotypic variation, we conducted single-marker regression GWAS. Overall, we detected 50 associated SNPs for all the traits, with 11 for 2018 and 19 for 2020 seasons, and 20 for the combined data that were spread along the entire sesame genome. The flowering date was mapped to two genomic regions (13 SNPs) on LGs 2 and 11 (Fig. 5A and Supplemental Table S9). The major genomic region on LG2 detected in the current study under the Mediterranean climate was previously reported in another sesame panel, however, it was less significant under other environments (Wei et al., 2015). The advantage of using a diversity panel to detect new alleles is exemplified by comparing our results with a bi-parental population that was grown under the same environmental conditions. While Teboul et al. (2020) detected six QTLs using the F_2_ population (S-91 × S-297), only the genomic region on LG11 was overlap with the current study.

The plant architecture traits were associated with 9 SNPs [HTFC (4), RZ (2), and RI (3)], with some overlaps (Fig. 5B, 5C; Supplemental Fig. S4A; Supplemental Table S9). Two genomic regions on LG 2 and 16 showed overlaps for HTFC and RI, which correspond to a high genomic correlation (r=-0.96) between those traits. These results suggest that genotypes with shorter HTFC will have the potential to extend the growth period and develop more capsules (i.e., high RI). Notably, while FD and PH showed positive phenotypic and genomic correlations (0.70 and 0.58, respectively), we did not detect any genomic region associated with PH. On the other hand, clustering analysis of the phenotypic data suggested that different genotypes with similar PH can differ in FD. Thus, the lack of genomic region associated with PH may result in either many small effect loci or management effect. It is also worth noting that since sesame is grown as a summer crop under irrigation management and characterized with indeterminate growth habits, it might affect the diversity of this trait (Supplemental Table S3).

Twenty-eight SNPs were significantly associated with yield components. Of those, 20 were associated with SYPP, 7 for SNPP, and 1 for TSW. Seed size (TSW) is known to have moderate-high heritability in sesame (Uzun et al., 2013; Kalaiyarasi et al., 2019), as was found in the current study (0.88) and other crops, such as wheat (Sukumaran et al., 2018) and pea (*Pisum sativum* L.; Huang et al., 2017), was associated with only one region in 2020, with no overlap with SNPP and SYPP. Likewise, a small number of associated loci for TSW were found in bi-parental sesame populations (Du et al., 2019; Teboul et al., 2020), which may suggest that this trait is under less complex genetic control or under the regulation of many small-effect genomic regions. The absence of significant genomic signs for TSW (as well as other traits in the current study) indicates that other approaches, such as genomic prediction (Crossa et al., 2017), may contribute to understanding its genetic basis. The positive phenotypic and genomic correlations between TSW and SYPP (0.37 and 0.44, respectively), and between SNPP and SYPP (0.86 and 0.93) on one side, and the absence of both correlations between TSW and SNPP (−0.04 and 0.17, respectively) on the other side, open up the possibility to breed simultaneously for both traits and improve yield.

SYPP and SNPP are highly polygenic traits associated with various anatomical and morphological traits (e.g., NBPP, RZ, RI, PH, number of capsules per plant, number of capsules per leaf axil, carpel number per capsule, and seed size). For SNPP and SYPP, we found one shared genomic region on LG2 overlapping with FD (Supplemental Table S9). This demonstrates the important role of FD on productivity, which is supported by the negative phenotypic and genomic correlations between these traits (Fig. 4). The effect of flowering on seed yield can be also attributed to the in-determine growth habit of sesame, and the agronomic practices to stop the irrigation to harvest the crop before it rains in autumn. Under such conditions, plants that were able to flower earlier had a longer period to produce flowers and capsules and obtain a higher yield.

In contrast, genotypes that flowered later in the season were more exposed to environmental conditions. A similar pattern was found in other in-determined crops such as soybean (Zhang et al., 2015) and chickpea (*Cicer arietinum* L.; Upadhyaya et al., 2015). It is worth noted that while FD had high broad-sense heritability (0.97), both SYPP and SNPP had relatively lower broad-sense heritability estimates (0.74 and 0.66, respectively) (Table 1). These results, along with haplotypes analysis (Fig. 7), suggest that these yield components are controlled by other less heritable factors not related to FD.

In sesame, allelic variation within flowering-related genes contributes to variation in flowering date (Wei et al., 2016; Zhou et al., 2018). In the current study, we found two genomic regions (LGs 2 and 11) that were associated with FD across years. The significant marker on LG11 is located within LOC105173174, which encodes to AT-hook motif nuclear-localized protein 9. The members of this gene family are associated with the regulation of flowering date in other plants species (Zhao et al., 2014). The co-localization of the major genomic region on LG2 for FD and yield components together may suggest that this region contains one major gene with a pleiotropic effect or cluster of several genes. Analysis of CGs within this genomic region highlighted 20 flowering and productivity-related genes (Supplemental Table S10). LOC105156148 is encoding nitrate transporter (NRT1) and was found in *Arabidopsis* to interact with two flowering regulators transcription factors, *CONSTANS* and *FLOWERING LOCUS C* (FLC) (Teng et al., 2019). LOC105156159 is encoding abscisic acid receptor *PYR1-like*, a mutant allele of this gene is found to be associated with growth and productivity in rice (Miao et al., 2018). Two of the significant markers within this genomic region were within the LOC105156152 gene that encodes *CHROMATIN REMODELING 19* regulating floral organs in *Arabidopsis* (Wu et al., 2012). The functional annotation of these genes and the co-localization of them in the same genomic region demonstrated how the variation in FD and SYPP could be genetically controlled together. Further investigation is needed to study how this genomic region (and the genes inside it) interact with other identified genomic regions for a deeper understanding of the genetic basis and mechanisms underlying FD and SYPP variations in sesame.

## 5. Conclusion for future perspective

While sesame is still mostly grown under a traditional cropping system, future sesame breeding targets should focus on improving yield and adaptively to more diverse environments. Thus, elucidating the genetic architecture controlling phenological, morphological, and yield components traits will aid in understanding selection criteria and better genetic-based breeding. Here we established a new sesame panel (SCHUJI) and explored its genetic variation for morpho-agronomic traits under the Mediterranean climate conditions. We showed the benefits of using the globally distributed panel for discovering new alleles associated with these traits. A major genomic region on LG2 was found in association with the flowering date and yield components, indicating the crucial role of phenology on sesame production. A better understanding of the genetic variability underlying flowering date in sesame will serve as a basis for improving sesame adaptively to new cropping systems. Moreover, it will enhance breeding efforts and enable turning this important crop from domestically grown to global production in intensive agriculture.

## Abbreviations

BLUE: Best linear unbiased estimate
CGs: candidate genes
FD: flowering date
GWAS: genome-wide association studies
HTFC: height to the first capsule
NBPP: number of branches per plants
PH: plant height
RI: reproductive index
RZ: reproductive zone
SNP: single nucleotide polymorphism
SNPP: seed number per plant
SYPP: seed yield per plant
TSW: thousand-seed weight

## ACKNOWLEDGEMENTS

We highly appreciate the excellent technical support of the Peleg lab members. This research was partly supported by the Chief Scientist of the Israel Ministry of Agriculture and Rural Development (837-0150-14), and The Hebrew University of Jerusalem.

## Notes

### Competing Interest Statement

The authors have declared no competing interest.

