## Supplementary material for "Genome-wide association uncovers the genetic architecture of tradeoff between flowering date and yield components in sesame": SI Figures

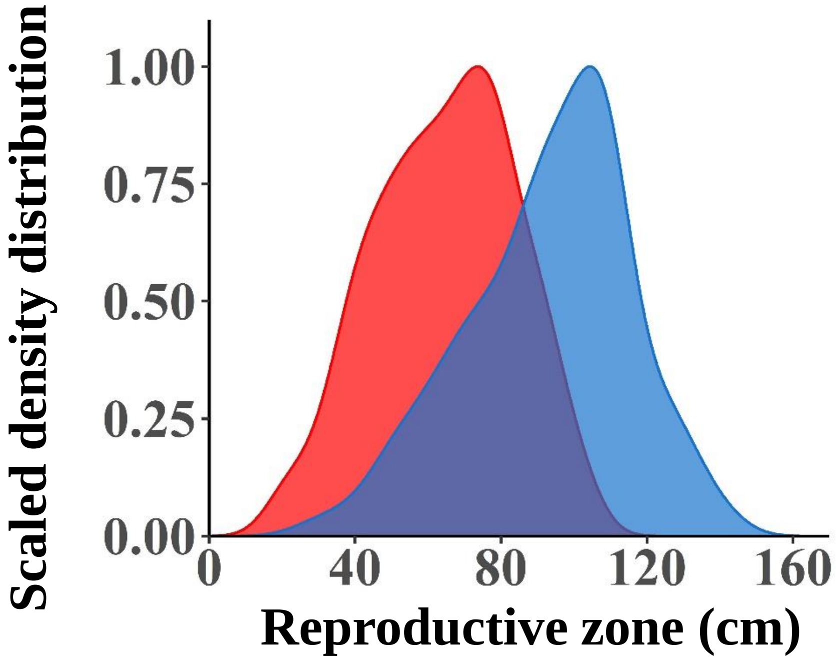


**Supplemental Figure S1**. Density distribution of reproductive zone.

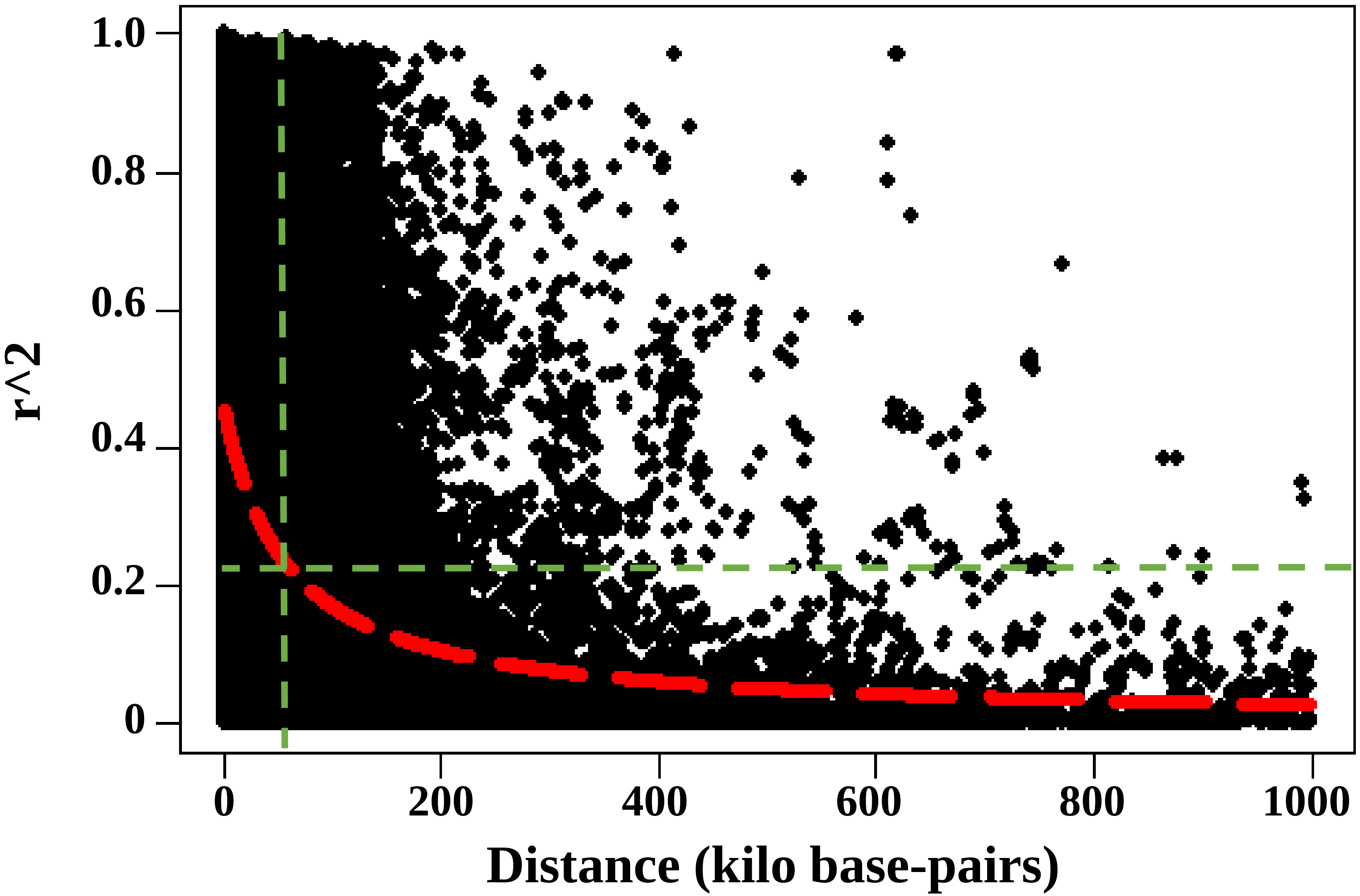


**Supplemental Figure S2.** Genome-wide linkage disequilibrium pattern in the SCHUJI panel. Red dashed line represents the non-linear trend and green dashed line indicates the point where linkage disequilibrium drops to half of its initial value (0.22) at 58,774 base pairs.

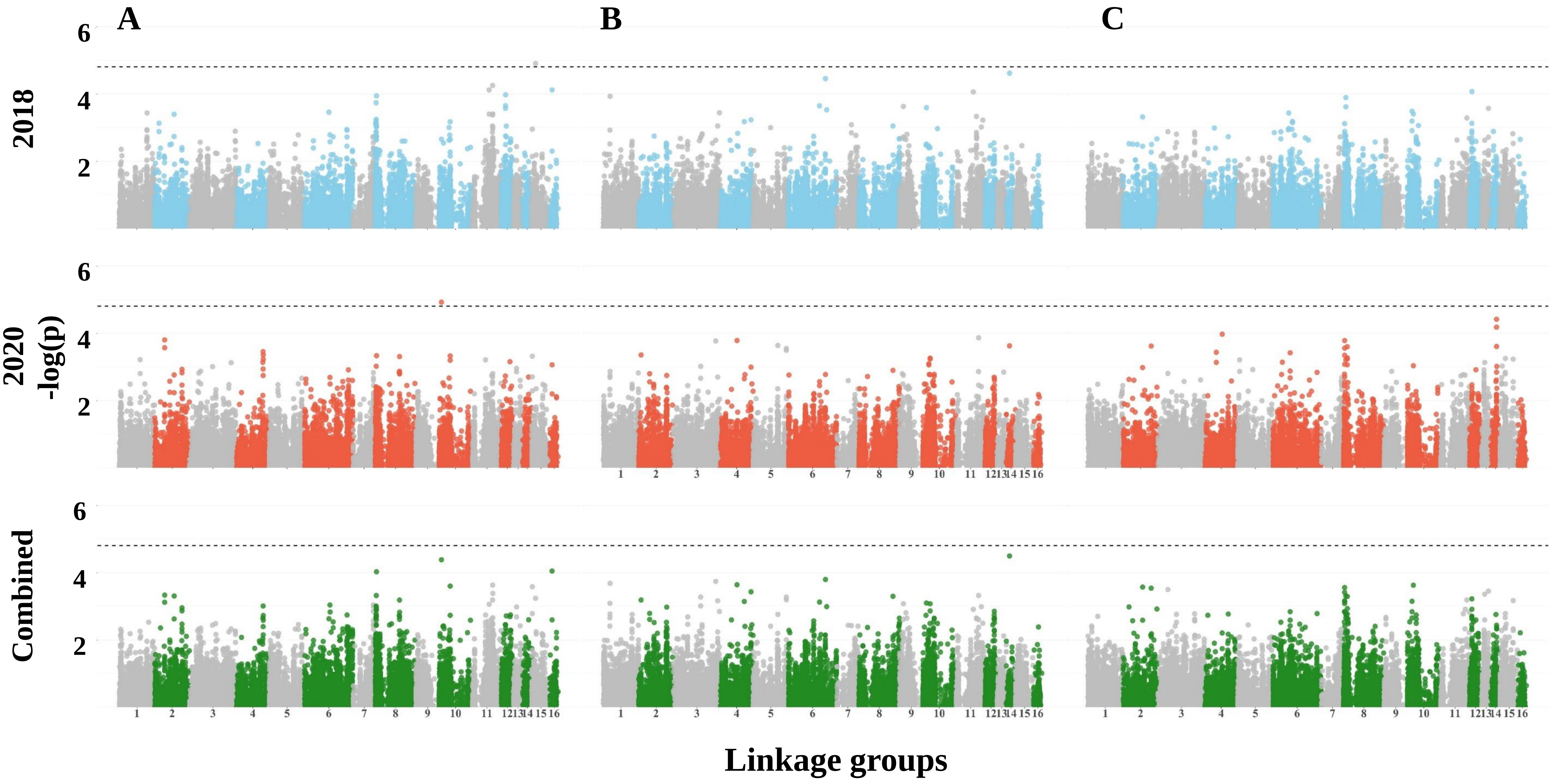


**Supplemental Figure S3.** Manhattan plots for (**A**) reproductive zone, (**B**) plant height, and (**C**) number of branches per plant. Dashed line represents the genome-wide significant threshold.

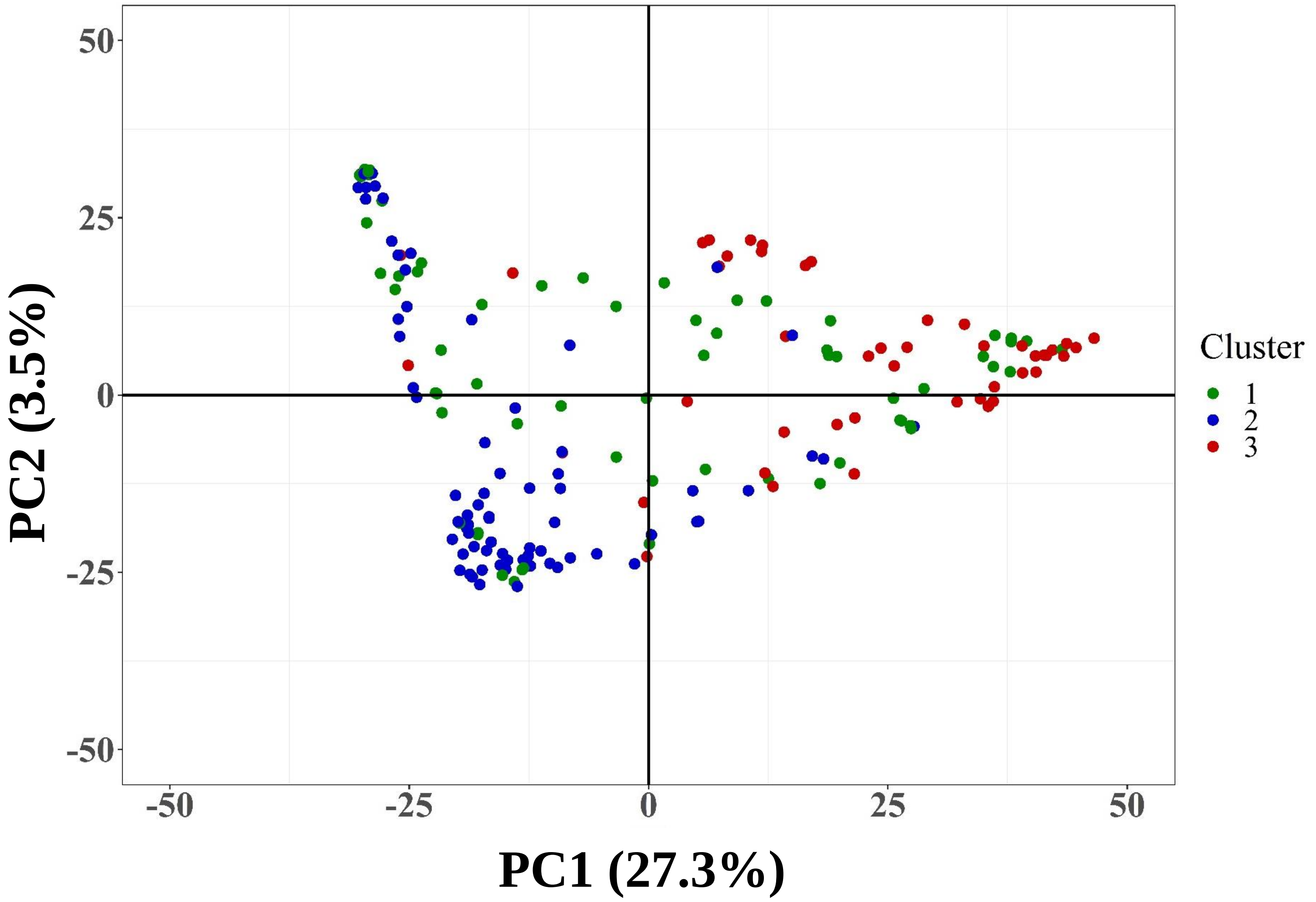


**Supplemental Figure S4.** Principal component analysis on genetic markers of the SCHUJI panel. Colors represent different cluster from the k-means cluster analysis. Red color represent cluster 3, green color represent cluster 1 and blue color represent cluster 2.
